# Pupil size variations reveal Bayesian inference in cognitive arithmetic

**DOI:** 10.1101/2024.11.27.625652

**Authors:** Alexandre Zénon, Samuel Salvaggio, Michael Andres

## Abstract

The assumption that the brain relies on Bayesian inference has been successful in accounting for many behavioural and neurophysiological observations, but to date, dependence on such mechanism has not been assessed in the context of arithmetic. Bayesian inference implies the representation of uncertainty and reliance on prior beliefs. In arithmetic problem solving, it would consist in refining prior knowledge about the response range as the system progressively integrates the numerical information conveyed by the operands. Within this framework, the amount of information needed to progress from a prior to a posterior probability distribution over responses can be quantified by the information gain, which would relate to the cognitive workload of the task. To test this hypothesis, we designed three experiments in which participants computed the sum of two numbers presented one after another through headphones. In each experiment, the information about response predictability conveyed by the first operand was manipulated. The first operand was either highly informative and contributed to narrow down the response range, or poorly informative and conveyed little information about a plausible response. Throughout all experiments, we found that pupil-related arousal signalled the information gain associated with the first operand, indicating that participants already updated the probability distribution of possible responses upon hearing that first stimulus. This finding shows that Bayesian inference is central to arithmetic problem solving and that information gains consecutive to the integration of the operands can be tracked over time through pupillometry.

**Public significance statement:**

– This study suggests that when we do arithmetic, our brain uses Bayesian inference, a method similar to how scientists update their theories based on new evidence. This means that when doing math, we start with hypotheses about the result and progressively narrow down possible solutions.
– By observing the eye of the participants while they were solving arithmetic problems, the authors showed that their pupil size increased in proportion to the amount of processed information.
– The authors also found that when participants received more informative first operands in the math problem, and when their pupil size increased more during this presentation, they were also faster to respond, indicating that greater initial information processing facilitated quicker problem-solving.

## Introduction

Bayesian inference is viewed as a core mechanism of the human brain, providing a comprehensive account of behavioural and neurophysiological observations (Colombo & Seriès, 2012). Reliance on Bayesian inference implies that individuals tackle problems with some *a priori* knowledge (or probability distribution) of the expected responses and update these beliefs upon reception of new pieces of information (Knill & Pouget, 2004). Bayesian inference has proved effective in accounting for a wide range of human skills, from perception (De Lange et al., 2018) to sensorimotor learning (Körding & Wolpert, 2004). A challenge for these probabilistic theories is to account for the computations underlying culturally inherited functions, such as language or mathematics, which require processing the meaning of symbols (Xu & Tenenbaum, 2007). Here, we question whether Bayesian inference can be considered a core mechanism for solving arithmetic problems. Reliance on prior distributions has been proposed to explain performance in addition (Ashcraft, 1987), multiplication (Siegler, 1988), as well as number comparison (Rinaldi et al., 2022). These models typically refer to statistical properties of arithmetic learning, such as frequency or order of presentation in textbooks, to account for the strength of problem representations in long-term memory (De Visscher & Noël, 2014; Hamann & Ashcraft, 1986). They share the assumption that arithmetic problems encountered more often are processed more efficiently (Ashcraft, 1992). Such processing advantage is compatible with the view that, through repeated practice, priors over responses have been adapted to fit frequent problems particularly well. However, these probabilistic models are concerned with overlearned problems represented in an organized network in memory (e.g., 2×4 = 8), and it is not clear how Bayesian inference can account for the solving of complex arithmetic problems (e.g., 43+8 = 51) whose difficulty lies in the need to process the value of the numbers and apply procedures.

We aim to demonstrate the inherently probabilistic nature of arithmetic problem solving by providing a mechanistic account of the relationship between arithmetic difficulty and information gain (IG), a measure of the divergence between prior and posterior beliefs that has recently been proposed as a universal currency to express cognitive cost (Zénon et al., 2019). This framework provides a theoretical basis for addressing the determinants of arithmetic difficulty and a quantitative assessment method leveraging the pupil response. To address the question of the reliance on prior beliefs for arithmetic problem solving, measuring RTs may not be sufficient as they index the final outcome of all serial or parallel processes involved in the task, reflecting the sum of all computations from the encoding of the stimuli to the solving of the arithmetic problem. However, updating of evidence occurs in multiple, successive steps: encoding the first operand (e.g. 47), the operator (e.g. + or –) and the second operand (e.g. 4) allows refining progressively the prior distribution of the probabilities assigned to different candidate responses within a plausible range. In other words, each operand contributes to reducing the uncertainty about the response and is thus associated with a specific and quantifiable IG. In order to assess the independent contribution of each operand to the cognitive cost of the solving of arithmetic problems, we chose to measure changes in pupil dilation from the presentation of the first operand, thus long before a final decision could be made on the response. Beside light, pupil size is influenced by several affective and cognitive factors such as emotions, surprise or mental effort (Alamia et al., 2019; Bradley et al., 2008; van der Wel & van Steenbergen, 2018), and was also found to increase consecutively to the solving of arithmetic problems (Hess & Polt, 1964). The ensemble of these findings can be explained by the hypothesis that pupil size reflects IG (Zénon, 2019), and we will use the pupillary signal as an index of this information theoretic measure of cognitive cost.

We conducted a series of experiments where the pupil size of adult participants was recorded while they solved addition problems with two-digit numbers presented sequentially in auditory format. The participants were informed about the range of all possible responses before the experiment, so that they all had the same prior internal models about the probable distribution of the responses. The information conveyed by the first operand was manipulated differently across conditions so that it could be more or less informative, thus more or less efficient in narrowing down the range of possible responses, to test the prediction that mental arithmetic works as a form of Bayesian inference (Figure 1).

**Figure 1.**
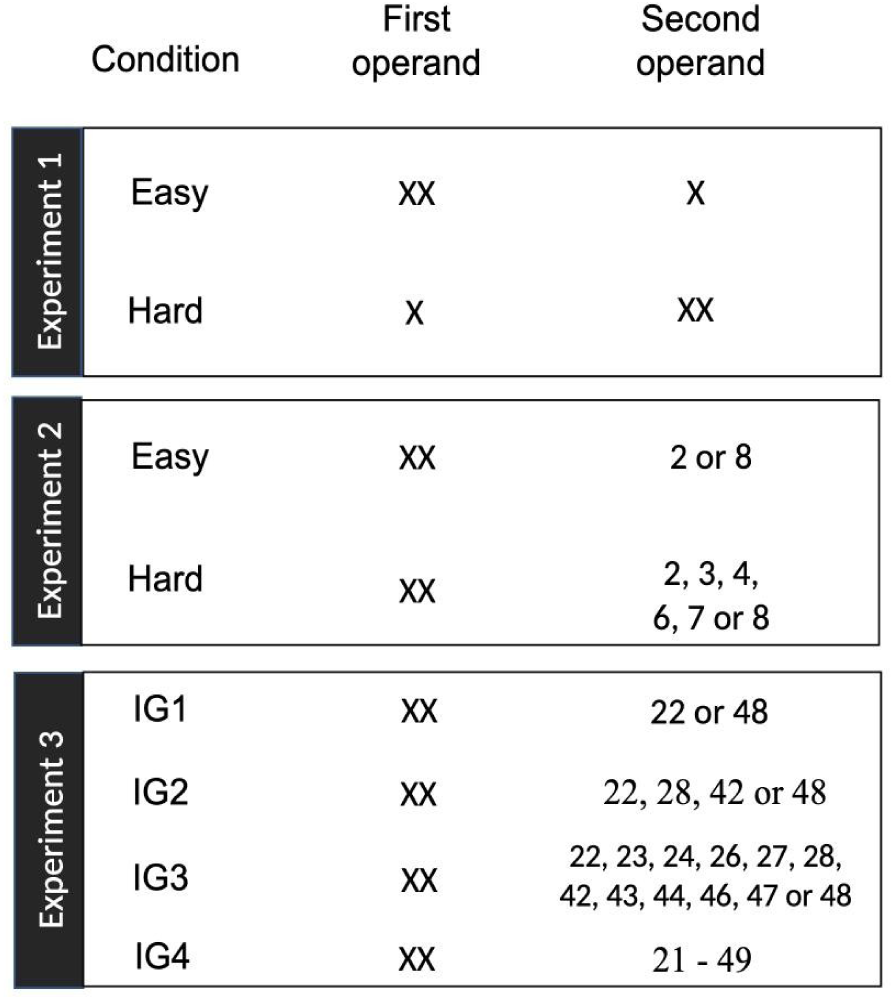
Schematic depiction of the three experiments. The task consisted of a mental addition task, with the two operands being presented successively and auditorily. The difference across experiments was the choice of values for those operands.

## Experiment 1

The first experiment used addition problems whose operands consisted of a two-digit number and a single digit. In such problems, the two-digit number was highly informative because it greatly reduced the range of responses, compared to the single digit which was less informative in that respect. The experiment was based on a within-subject design, with two sessions consisting of the same addition problems, except that the order of the operands was reversed between sessions to manipulate the information gain associated with the first operand. In one session, the first operand was the two-digit number and the second operand was the single digit (XX + X). In the other session, the first operand was the single digit and the second was the two-digit number (X + XX). We predicted that, after hearing the first operand, pupil dilation should vary as a function of the information gain. Specifically, given the format of the addition problems tested, pupil size should dilate more when the first operand is a highly informative two-digit number than when it is a single digit with low informative value.

## Method

### Participants

Twenty French-speaking participants took part in this experiment (nine males; two left-handed; mean age and standard deviation, *M*=20.7, *SD*=1.8). They all had normal or corrected-by-lenses vision and did not report any antecedent of mathematical learning disability when asked by the experimenter. The experiment was in accordance with the ethical standards established by the Declaration of Helsinki and was given prior approval by the local ethics committee. Due to the lack of literature on the effect we sought to test, the sample size was arbitrarily preplanned to have a balanced and sufficient number of participants for the various experimental conditions and randomisations.

The study complies with APA ethical standards in the treatment of the acquired data.

### Apparatus

The experiment was run on a PC equipped with a 22-inch LCD screen (1920×1080 pixels; refresh rate: 60Hz). The Matlab software (v.2018a) controlled the stimulus presentation and the recording of the verbal response (MathWorks Inc., 2018). An Eyelink 1000 tower-mounted camera was used to track eye movements (SR Resarch, Mississauga, Canada; sampling rate: 1000Hz; average accuracy range: 0.25° angle to 0.5° angle; gaze tracking range of 32° angle horizontally and 25° angle vertically). Participants were placed at 60 cm from the screen. Prior to each experimental block, the eye tracker was calibrated to the screen using a built-in 9-point protocol.

### Stimulus materials

The auditory stimuli consisted in either double digits French number words whose magnitude ranged from 21 to 69, excluding numbers ending in “0” (i.e., 30, 40, 50, 60) or single digits numbers from “1” to “9”. The number words were recorded in a stereo audio file whose duration was adjusted to 750ms for two-digit numbers and 500ms for one-digit numbers. The screen stayed unchanged for the duration of all the experiment. It consisted of a central white fixation dot on a grey background. The experiment counted 144 trials for each condition, giving 288 trials in four separated blocks. Thus, there were two blocks where the stimulus order consisted in a double digit followed by one digit and two blocks where the stimulus order was single digit followed by double digit. We used the Kullback-Leibler (KL) divergence between prior and posterior beliefs to quantify the IG associated with the first operand both in the double-digits first blocks and in the single-digit first blocks. The KL divergence can be calculated, for probability distribution *P* and *Q*:

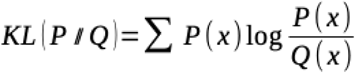

In the present experiment, the KL divergence of the two-digit number first, or high informative condition was 2.38 bits while the KL divergence of the one-digit number first, or low informative condition was 0.24 bits.

### Task and Procedure

Each trial was preceded by a 1000ms fixation cross. The trial started with a number word that was played in the headphones while the screen stayed unchanged. Then, 2750ms after trial onset, the second number was played in the headphones. Participants had to add the two numbers and answer orally. The display remained in this configuration for 3,750 additional ms while participants’ pupil diameter was still being recorded. The next fixation cross started immediately after. Participants were also instructed to look constantly at the fixation cross that remained in the centre of the screen during all the experiment. Before starting all block of trials, they were reminded of the range of the possible response and the order of stimuli (i.e. double digit or single digit first). The order of the blocks was counterbalanced across all participants.

### Data analysis

Erroneous trials were excluded from analyses (10.3% of dataset). Response times (RTs) corresponded to the delay between the onset of the second operand and the onset of the subject’s response. Pupillary data was pre-processed by linearly interpolating missing data resulting from eye blinks and by high-pass filtering it (digital Chebyshev filter, 1/16 Hz cutoff frequency). We then isolated the impulse response of the pupil to cognitive events (operand presentations and response) by running an ARX analysis (Zénon, 2017). The orders of the ARX model were fixed to 3 (autoregressive process) and 15 (exogenous inputs) whereas the delay parameter was chosen within a range of 1 to 5, so as to maximise the peak of the impulse response. The pupillary response to the operand presentations and the response in each trial was then obtained by running a GLM with the pupil signal as explained variable and three regressors per trial, each obtained from the convolution of the event vectors (zero everywhere but at the time of occurrence of the events) with the subject-specific, pupillary impulse response. This resulted in one regression coefficient per event per trial, which represented the magnitude of the response to that event at that trial. The autocorrelation in the residuals was taken care of with an AR-2 model (Cochrane & Orcutt, 1949). We then ran Generalized Linear Mixed Models (GLMM) with the coefficients corresponding to the first operand as dependent variable and IG condition (2.38 bits vs. 0.24 bits) as independent variable: *Pupil size ∼ 1 + IG + (1+IG |Participant).* The dependent variable was transformed so as to make its distribution closer to normal:

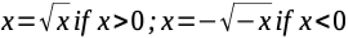

Residuals were systematically inspected by means of qq-plots and per-participant boxplots. Degrees of freedom were estimated with Satterthwaite’s approach and estimation of model parameters was performed with REML method. We also ran a GLMM on log-transformed RTs with IG condition and z-scored pupil dilation as dependent variables: *RTs ∼ 1 + Pupil + IG + (1+pupil+IG |Participant)*.

All analyses were conducted with MATLAB 2018a (MathWorks Inc., 2018) and Jamovi (The jamovi project (2021). *jamovi* (Version 1.6) [Computer Software]. Retrieved from https://www.jamovi.org).

### Transparency and openness

We report how we determined our sample size, all data exclusions (if any), all manipulations, and all measures in the study. All data and statistical models are available at https://researchbox.org/3762. Data were analyzed using Matlab version R2016b (The Mathworks Inc.; code available at https://github.com/alexandre-zenon) and Jamovi version 2.5.4.0 (The Jamovi project).

## Results

We found that the IG condition affected pupil dilation significantly, with larger pupil responses being associated with larger IG (*F*_(1,19.2)_=14.3, *p*=0.001, *β mean and [95% CI]:* 0.155 [0.0744,0.2348]; see Figure 2A), in agreement with our hypothesis. The RTs were found to depend also on condition (*F*_(1,19.1)_=30.6, *p*<0.001, *β mean and [95% CI]: –0.2452 [-0.332,-0.1583]*), with longer RTs in trials with large IG, though these conditions corresponded to trials with double-digit operands, whose presentation lasted 250ms longer. This makes it difficult to determine whether the effect of condition on RTs was due to information content or to the duration of the presentation. However, and more importantly, we found that larger pupil dilation to the first operand led to shorter RTs (*F*_(1,5279.7)_=76.5, *p*<0.001, *β mean and [95% CI]: –0.2654 [-0.325,-0.206]*; see Figure 2A), confirming that the amount of information being processed during first operand presentation was instrumental in decreasing computation time at the moment of second operand presentation.

**Figure 2.**
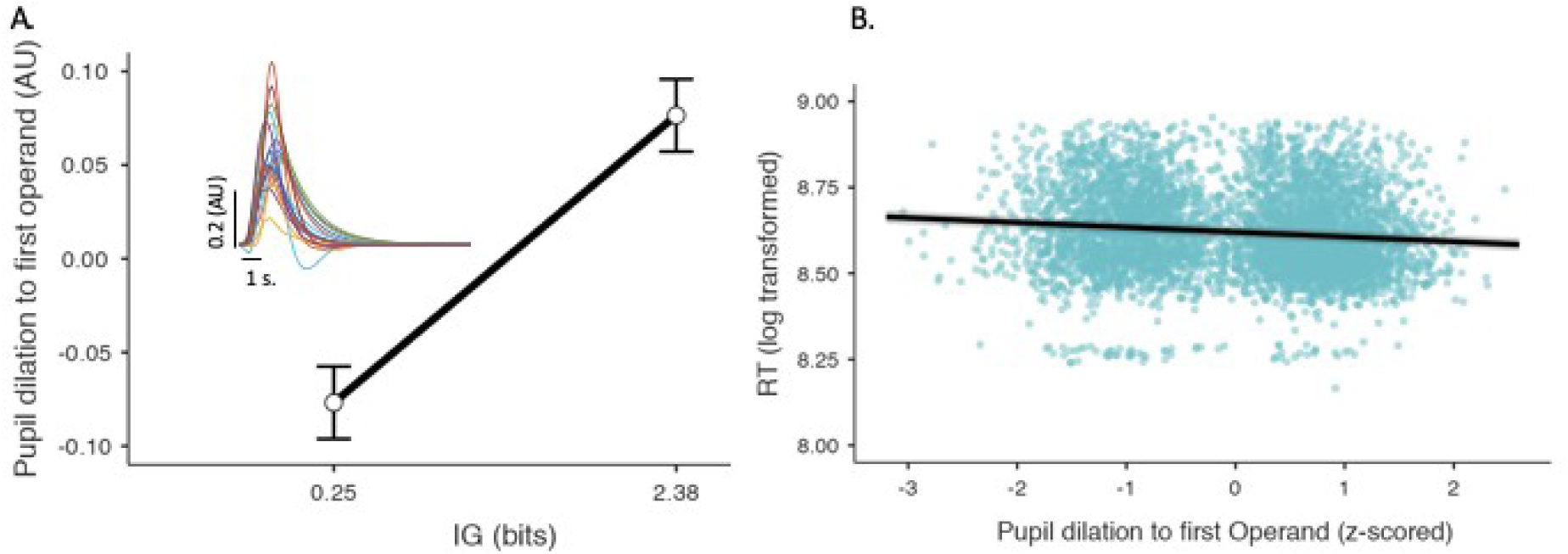
A. Average pupil dilation to first operand presentation as a function of its information gain in bits. Error bars indicate standard error of the mean. Insets in the upper left corner shows individual impulse responses to operand onset. B. RT (log transformed) as a function of pupil dilation to first operand. Confidence interval of the slope is indicated in gray shade. Each dot corresponds to an individual trial.

## Interim discussion

In this experiment, participants’ pupil size was recorded while they were solving addition problems. Each operand was presented sequentially and the amount of information provided by the first operand was directly manipulated. Results showed that the pupil response to the presentation of the first operand was significantly higher in the 2.38 bits condition when compared to the 0.24 bits condition. The analysis of RTs showed that problems where the first operand was less informative were also more difficult and took longer to be resolved. We also highlighted a negative relationship between average pupil size and RT. These results converge to show that pupil dilated in relation to the IG associated with the first operand, which helped participants to narrow down the range of possible solutions beforehand, helping them to solve the arithmetic problem faster. They are consistent with the hypothesis that arithmetic problem solving is based on probabilistic inference and relies on progressive updating of prior distributions over possible responses, as new information is received.

Past research has long been animated by the question of which problems are solved by procedural strategies and which are resolved through memory retrieval strategies (Ashcraft, 1992; Chen & Campbell, 2018; Fayol & Thevenot, 2012). Procedural strategies, such as counting or decomposition, were initially associated with longer response latencies and higher error rates, compared to retrieval strategies, underlining the role of arithmetic difficulty in the selection of the solving strategy (Campbell & Xue, 2001). However, previous studies mainly investigated the solving of single-digit arithmetic problems (Nuerk et al., 2011). In comparison, little is known about the difficulty of two-digit arithmetic problem solving. Conventionally, difficulty indicators consist of categorial criteria, such as the total number of digits or the need to carry over digits across the tens places, which have been used to account for increased RTs and error rates (FÜrst & Hitch, 2000; Klein et al., 2010; Moeller et al., 2011; Seitz & Schumann-Hengsteler, 2002). However, these indicators are limited since they fail to provide an operational definition of arithmetic difficulty for all problems – only for categories – and by extension they fail to provide a comprehensive account of the cognitive cost of arithmetic problem solving at a more conceptual level. The present results provide a complementary account of two-digit arithmetic difficulty, based on the progressive narrowing down of the range of solutions, quantified through information gain.

However, the use of single digits or double-digit numbers as first operands raises alternative interpretations to the effect found on the pupil size. Indeed, the encoding of the first operands differed in phonological and lexical complexity between the two lists and the auditory presentation of a two-digit number was 250 ms longer than the auditory presentation of a single digit, resulting in a higher working memory load in trials involving two-digit numbers. We therefore conducted a second experiment where the first operands were the same but their informative value changed as a function of the second operands used in each list of problems, with the aim to replicate the effect of the information conveyed by the first operand on pupil dilation while excluding other sources of arousal increase.

## Experiment 2

To counteract possible biases in pupil size measures due to phonological or lexical differences related to the hearing of single-digit versus two-digit numbers as first operands, we conducted a second experiment where the first operand was always a two-digit number taken from the same list in both the low and the high information gain conditions. The informative value of the first operand in these two conditions was actually given by the relative value of the second operand: in the high IG condition, the second operand was chosen in a list of two possible digits, whereas in the low IG condition it was chosen in a list of six possible digits, leaving more uncertainty about the arithmetic outcome. We predicted a larger pupil dilation after hearing the first operand in the high information gain condition where the second operand carried little additional information about the outcome of the calculation.

## Method

### Participants

Twenty French-speaking participants took part in this experiment (eight males; one left-handed; mean age and standard deviation, *M*=22.2, *SD*=3.2). They all had normal or corrected-by-lenses vision and did not report any antecedent of mathematical learning disability when asked by the experimenter. The experiment was in accordance with the ethical standards established by the Declaration of Helsinki and was pre-approved by the local ethics committee.

### Stimulus materials

The first operands consisted in double-digit French number words whose magnitude ranged from 21 to 49, excluding numbers ending in “0” (i.e., 20, 30, 40). The second operands consisted in either {“2”, “8”} in the high information gain condition, or {“2”, “3”, “4”, “6”, “7”, 8”} in the low information gain condition. The experiment counted 162 trials for each condition and location of bright/dark part of the screen, giving 324 trials in six separated blocks. In the present experiment, the KL divergence of the high IG condition was 2.81 bits while the KL divergence of the low IG condition was 1.7 bits.

### Task and Procedure

The procedure was the same as in Experiment 1. Before starting all blocks of trials, participants were reminded of the range of the possible responses and of the second operand. The order of the blocks was counterbalanced across all participants.

### Data analysis

Two subjects were excluded from the analyses because of a large number of trials (>100) with poor pupil signal quality. Remaining trials with poor signal quality, as well as trials in which participants responded incorrectly were excluded from the analysis (10.9% of dataset). All analyses were conducted in a similar way to Experiment 1.

## Results

Similarly to Experiment 1, we found that pupil response to the first operand was larger under conditions of large IG (*F*_(1,17.4)_=5.17, *p*=0.036, *β mean and [95% CI]: 0.0723 [0.0099,0.1346]*; see Figure 3A). Likewise, RTs were also affected by IG condition (*F*_(1,16.9)_=22.6, *p*<0.001, *β mean and [95% CI]: –0.2132 [-0.301,-0.125]*) and pupil response to first operand (*F*_(1,17.2)_=31.9, *p*<0.001, *β mean and [95% CI]: –0.1038 [-0.140,-0.0677]*; see Figure 3B).

**Figure 3.**
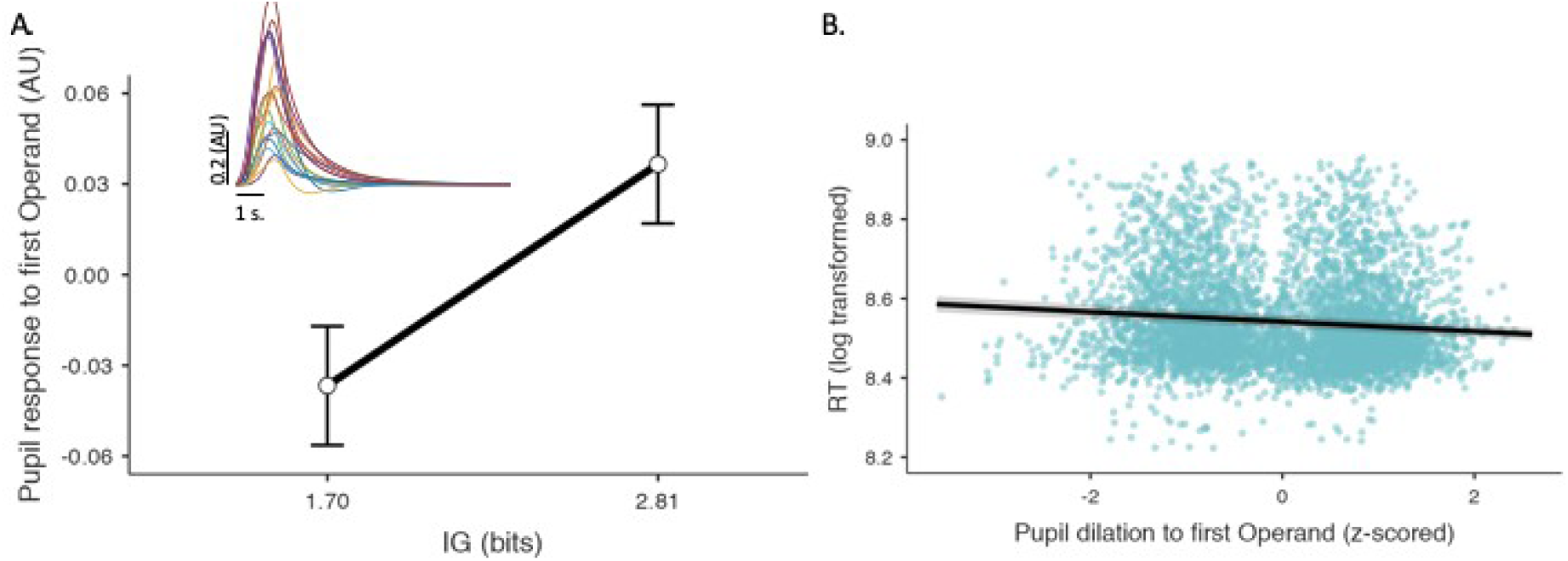
A. Pupil dilation to first operand presentation as a function of its information gain. B. RT as a function of pupil dilation to first operand. Same conventions as in Figure 2.

## Interim discussion

The results from Experiment 2 showed that, similarly to Experiment 1, in conditions in which the first operand was more informative, the eye pupil dilated significantly more and was associated with shorter RTs. These findings confirm our initial interpretation that hearing the first operand leads to updating of the priors over possible responses, in agreement with a Bayesian take on arithmetic problem solving. In the present experiment, the first operands were identical across conditions, meaning that the effect of the first operand on pupil dilation cannot be explained by low-level phonological or lexical differences.

However, Experiment 2 left open an alternative interpretation in terms of anticipatory calculation. Indeed, the rationale of this experiment was to match easy and hard conditions for the first operands but to make them more or less informative by varying the range of possible values taken by the second operands. Because the easy condition allowed only two possible values (+2 or +8), participants may have anticipated the result by computing both alternatives in parallel immediately after hearing the first operand. This strategy would provide them with a processing advantage that could explain the increase in pupil dilation in response to the first operand, compared to the hard condition, regardless of the associated IG. Experiment 3 was therefore motivated by the need to further specify the relationship between the information gain and the pupil response to the first operand. We created four different conditions that only differed in the number of possible values that the second operand could take: either 2, 4, 12 or 27 values. This allowed us to look for a linear relationship between pupil dilation and information gain across four different levels, including two additional conditions where anticipatory calculation was unlikely because it would require computing up to 12 or 27 additions in parallel.

## Experiment 3

In the third experiment, we created four independent conditions in which the first operand was always taken from the same list of two-digit numbers, whereas the second operand was a two-digit number taken from a list of varying length (i.e. 2, 4, 12 or 27 choices), resulting in four different levels of information gain. We expected a linear effect of information gain on the pupil size dilation, as measured after hearing the first operand, in the four different conditions.

## Method

### Participants

Twenty French-speaking participants took part in this experiment (eight males; one left-handed; mean age and standard deviation, *M*=23.4, *SD*=3). They all had normal or corrected-by-lenses vision and did not report any antecedent of mathematical learning disability when asked by the experimenter. The experiment was in accordance with the ethical standards established by the Declaration of Helsinki and was approved by the local ethics committee.

### Stimulus materials

The first operands consisted in double-digit French number words whose magnitude ranged from 21 to 49, excluding numbers ending in “0” (i.e., 20, 30, 40). The second operands were separated in four different condition. In the first, most informative condition, they were {“22”, “48”}. In the second most informative condition they were {“22”, “28”, “42”, “48”}. In the third condition, they were {“22”, “23”, “24”, “26”, “27”, “28”, “42”, “43”, “44”, “46”,”47”,”48”}. In the fourth condition, they could consist in any number between 21 and 49, with the exclusion of numbers ending in “0”. The experiment counted 108 trials for each condition, giving 432 trials in eight separated blocks. In the present experiment, the IG was 3.22 bits for the first condition, 2.53 bits for the second, 1.44 bits for the third, and 0.59 bits for the fourth.

### Task and Procedure

The procedure was the same as in Experiment 1 and 2 except for the longer fixation duration (7 seconds). This was added in order to further ensure that conditions would not differ in tonic pupil size. The experiment was divided in two different sessions in order to avoid participant fatigue. Each session had one k of each condition. Before every block of trials, participants were reminded of the range of the second operand and of the possible responses. The order of the blocks was counterbalanced across all participants and this order was kept constant between the two sessions.

### Data analysis

Three sessions from different participants were excluded from the analysis due to the poor quality of the pupil signal (more than 100 trials without usable signal). All analyses were performed similarly to previous Experiments, except that the IG associated with the four conditions of arithmetic difficulty was introduced as a continuous variable.

## Results

The analysis revealed a significant effect of IG on pupil dilation to first operand (*F*_(1,18.4)_=4.68, *p*=0.04, *β mean and [95% CI]: 0.0559 [0.0052,0.1066]*). Similarly to the other experiments, the RT analysis highlighted a significant main effect of IG (*F*_(1,19.4)_=8.81, *p=*0.008, *β mean and [95% CI]: – 0.0875 [-0.1452,-0.0297]*) and pupil size (*F*_(1,17.3)_=12.98, *p*=0.002, *β mean and [95% CI]: –0.0572 [-0.0884,-0.0261]*; see Figure 4B). In order to further support the hypothesis of a linear relationship between IG and pupil dilation, we aggregated the data of the three experiments and replicated the analysis on pupil dilation with IG as a continuous variable (Figure 4C). The results confirmed the linear effect of the IG used across the three experiments on pupil dilation to the first operand (using both subjects and experiments as random factors: F(1,58)=14.5, p<0.001, *β mean and [95% CI]: 0.1687 [0.0811,0.256]*).

**Figure 4.**
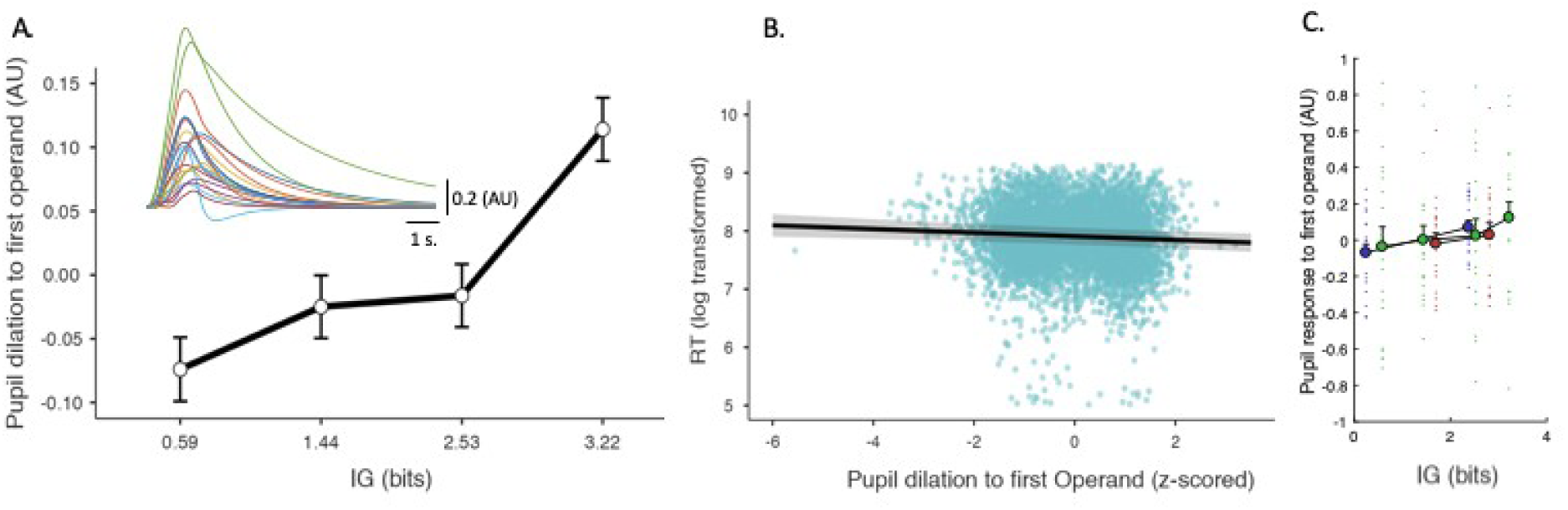
A. Pupil dilation to first operand presentation as a function of its information gain. B. RT as a function of pupil dilation to first operand. Conventions are identical to those of Figure 2. C. Pupil dilation as a function of information gain across experiments. Individual dots correspond to individual subjects.

## General discussion

The present study aimed to investigate pupil dilation in relation to the updating of prior distributions over possible responses in arithmetic problem solving. Across the three experiments, we found that the first operand induced a larger pupil dilation in conditions where it carried more information. Moreover, in all experiments, this increase of pupil size was negatively correlated with RT, showing that the informational value of the first operand, reflected in pupil dilation, helped participants solve the arithmetic problem. These findings corroborate the hypothesis that arithmetic problem solving relies on progressive updating of prior distribution over possible responses, in agreement with Bayesian models of behaviour (Colombo & Seriès, 2012; Penny, 2012) and that information gain can be used as measures of the cognitive cost of arithmetic problem solving and is a decisive factor of its perceived difficulty (Zénon et al., 2019).

This conclusion does not contradict previous accounts of arithmetic problem solving, that have highlighted the effect of problem size (Campbell, 2005; Verguts & Fias, 2005) or carry operations on associated workload (FÜrst & Hitch, 2000). On the contrary, the perspective we defend in this work could help explain some of these effects. The problem size effect refers to the decreased performance observed when individuals have to process larger operands in arithmetic problems, and has classically been attributed to frequency effects, larger numbers being encountered less frequently than smaller ones (Campbell, 2005). The Bayesian perspective used here is in agreement with this view, since more frequent encounters of the operands, together with their sum and difference, would favour their representation in the prior probability distributions, thereby improving performance (Zénon et al., 2019). On the other hand, the carry effect refers to the finding that performance decreases with the number and the value of the carries that need to be maintained and added in the leftward column in multi-digit calculation (e.g. 246+381). The working memory load incurred by carry operations is not so high, as evidenced by the low impact of articulatory suppression, an experimental manipulation that prevents verbal rehearsal (FÜrst & Hitch, 2000; Imbo et al., 2007). Yet, executive interference has been observed when the calculation imposes a sequence of carries of increasing value. Such interference has been explained by the competition between two task sequences, one without and one with the execution of a carry (Imbo et al., 2007). This explanation can be subsumed under the Bayesian perspective proposed here, considering that adding a previously stored value imposes significant updating of the prior distribution over possible responses.

The grounding of arithmetic problem solving in an information processing framework also provides a new perspective on the interaction between exact and approximate arithmetic processes (Dehaene et al., 1999; Hyde et al., 2014; Park & Brannon, 2014; Sekeris et al., 2021). We propose that approximate arithmetic would allow individuals to perform an initial, fast shaping of the prior distributions over responses, which would then be used as a starting point for exact arithmetic computations. Such combinations of approximate and precise processes has been shown to be advantageous from the information theoretic perspective (Genewein et al., 2015). From a more mechanical point of view, this initial, rapid but approximate shaping of prior probability distributions over possible responses could reduce the complexity of the working memory manipulations by eliminating unlikely options. We recently found a trace of this heuristic process in the eye movements performed by adult and children solving addition and subtraction problems (Masson et al., 2024; Salvaggio et al., 2022). We used lateralized eye movements to test the hypothesis that addition and subtraction shift attention in opposite directions along a left-to-right oriented continuum representing numbers in ascending order. A time course analysis revealed that addition deviated the gaze rightward, compared to subtraction, as soon as participants had processed the first operand and the operator. The predictive nature of these attention shifts, which typically occur before the presentation of the second operand, clearly shows that their role is not to identify the exact answer but to give an approximative idea of plausible answers that would then be compared to the solution obtained by an exact procedure. We proposed a probabilistic model where attention shifts are embedded in working memory processes: spatial resources would be used to adjust the range of plausible answers as the calculation progresses, thereby reducing the part of the number sequence maintained in verbal working memory for precise incrementation. The present study provides direct empirical support to this probabilistic model of arithmetic problem solving, highlighting the need to consider prediction as a core determinant of the interactions between space and number (Hawes & Ansari, 2020).

In summary, the present work proposes a new viewpoint on mental arithmetic processes and their associated cognitive cost, based on the principle of Bayesian inference. Empirical validation of the proposed framework shows that Bayesian inference provides a theoretically valid explanation of human behaviour, which is not limited to basic sensorimotor functions but encompasses more advanced cognitive functions learned through cultural practice, such as symbolic arithmetic. At a more mechanistic level, our probabilistic model explains how approximate arithmetic can help refine the range of plausible responses in advance of defining exact arithmetic procedures, thereby alleviating the working memory load. Future research is needed to enlarge this framework to other dimensions of numerical cognition.

## Acknowledgements

AZ is a research associate at the Centre National pour la Recherche Scientifique (CNRS, France). MA is a research associate at the Fonds National de la Recherche Scientifique (FRS-FNRS, Belgium). The research was supported by grant T.0006.20 from the FRS-FNRS and grant ARC21/26-112 from the Fédération Wallonie Bruxelles (FWB, Belgium) as well as ANR VICONTE 2018-0524 and ANR CoCogIT 2019-0054.

